# G-quadruplex topologies determine the functional outcome of guanine-rich bioactive oligonucleotides

**DOI:** 10.1101/2024.12.01.626249

**Authors:** Prakash Kharel, Nupur Bhatter, Safiyah Zubair, Shawn Lyons, Paul J. Anderson, Pavel Ivanov

## Abstract

Guanine-rich nucleic acid sequences can exert sequence and/or structure specific activities to influence biological and pathobiological cellular processes. As such, it has been reported that different G-rich oligonucleotides (both DNA and RNA) can have cytotoxic as well as cytoprotective effects to the cells. However, the mechanisms of such a biological outcome are unclear. In this report, we report that G-rich DNA oligonucleotides (ODNs) that can form four stranded secondary structures called G-quadruplexes (G4s) can have topology-dependent biological outcome. Using different biochemical, biophysical, and cellular approaches, we demonstrate that only the parallel topology G4-forming ODNs can repress eukaryotic translation by directly interacting with eukaryotic translation initiation protein 1 (EIF4G1), while the antiparallel topology G4s do not have inhibitory effect on mRNA translation To the best of our knowledge, this is the first report to directly connect the G4 topological differences with differential functional biological impacts. Our study provides the foundation for the rational design of G-rich oligonucleotides for a desired therapeutic outcome.

## INTRODUCTION

Oligonucleotide based therapeutic molecules can regulate target gene expression via several different mechanisms such as RNA interference, target degradation by RNase H-mediated cleavage, splicing modulation, gene editing, and gene activation^1^. Among different categories of therapeutic oligonucleotides, guanine (G)-rich sequences bring an additional dimension due to their unique structural and functional properties^2,3^. G-rich oligodeoxynucleotides (ODNs) and oligonucleotides (ONs) have been reported to exhibit a wide range of biological effects, encompassing cytotoxic^4,5^, anti-cell proliferative^6^, and cytoprotective activities^7^. The antiproliferative and pro-apoptotic effects of G-rich ODNs/ONs have been observed in various cell types, including vascular smooth muscle cells and numerous cell lines derived from solid tumors, leukemias and lymphomas. Importantly, while most common nucleic acids based therapeutic strategies depend upon their anti-sense properties (complementarity of the drug-target pair), G-rich ODNs/ONs are known to exhibit unique non-antisense (for example, specific protein binding) properties when delivered to target cells^8^. The non-antisense antiviral effects of G-rich ODNs were first noted almost three decades ago^9^.

As such so-called G-rich T-oligos (DNA oligos homologous to the telomere 3′ overhang region, for example, pGTTAGGGTTAG) have been extensively studied as potent cytoprotective agents in normal cells as evidenced by their ability to induce transient cell cycle arrest in normal human cells of many lineages^5^. Importantly, G-rich T-oligos were shown to induce selective cytotoxicity on malignant cells compared to their normal counterparts and so they have been long touted as potent anti-cancer drugs^10,11^. For some ODNs/ONs, *in vivo* antiproliferative effects were also demonstrated in tumor-bearing mice. Furthermore, G-rich ODNs/ONs containing several backbone modifications can bring a diverse array of biological effects upon their cellular delivery, for example, inhibition of cell proliferation, induction of cell death, changes in cellular adhesion, inhibition of protein aggregation, and antiviral activity^12^.

Several bioactive G-rich ODNs/ONs can fold into four stranded secondary structures called G-quadruplexes (G4s) via stacking of square planar G-quartets^6,13^. G4s in both DNA and RNA possess a wide range of biological functions depending upon the context of their presence and their surrounding environment^13^. Several factors such as conformational polymorphism, loop permutations^14^, spatial restriction^15^, stability variations^16^ endow G4s with diverse functional applications as aptamers, fluorescence probes and nanodevices, biosensors and drugs^17^. In some cases, G4 ODNs and ONs facilitate their function by directly interacting with G4-binding cellular proteins (G4BPs)^18^ thus leading to cytotoxicity^6,19^ or cytoprotective^7^ effects. Alternatively, G-rich ODNs/ONs can bind to specific cellular proteins independent of G4 secondary structure^20^ to alter cell proliferation.

While most of the reported cellular G4s act *in cis*, the G4s formed by smaller nucleic acids including therapeutic ODNs/ONs mostly act *in trans* and obviously will have multiple targets. Even though G-rich ODNs/ONs have been reported to demonstrate unique, biological properties, it has been overlooked what makes some of the G-rich ODNs/ONs act differently than the others? In this study we analyzed the properties of several G-rich ODNs to induce several synergistic biological functions within stress responses and discovered that such ODNs are translationally active (active translation repressors) only when they fold into a parallel topology G4s (p-G4s). Importantly, our results demonstrate that a switch in G4 conformation from anti-parallel G4 (ap-G4) to p-G4 can revert their biological functions. Additionally, we identify molecular mechanism by which G-rich ODNs impact cell biology by targeting the global translation machinery in the human cells.

## RESULTS

### G-rich ODNs differ in their ability to induce mRNA translation repression

Translation of messenger RNA (mRNA) in eukaryotic cells is strictly regulated to modulate protein synthesis during cellular homeostasis, development, and cellular stress^21^. *In vitro* translation (IVT) systems based on rabbit reticulocyte lysate (RRL) have been extensively used to study the mechanism and outputs of translation *in vitro*^22^.Previously, we demonstrated that several tRNA-derived stress-induced RNA molecules (tiRNAs) can repress eukaryotic translation^23-25^. Importantly, two of the translation repressing tiRNAs contain 5’-**T**erminal **O**ligo**G**uanine (5’TOG) motif, which was shown to be responsible for the repressing activity of those particular tiRNAs ^18,23,26^. Furthermore, we and others demonstrated that several G-rich ODNs and ONs can impart different physiological outcomes in normal and stressed cells^7,27^. Because we previously showed that several G-rich RNAs can repress RRL-based *in vitro* translation of uncapped firefly luciferase mRNA in RRL^23^, we asked whether translation repression is the characteristic of all G-rich oligonucleotides regardless of their sugar composition. To answer this question, as schematically demonstrated in **Figure 1A**, we performed IVT experiments in the presence of five G-rich sequences (3 G-rich ODNs and 2 G-rich ONs) in addition to two control non-G rich ODNs and two positive controls (namely 5’tiRNA^Ala^ and 5’tiDNA^Ala^) in RRL. To our surprise, we observed that only 3 of test candidate G-rich ONs/ODNs repressed *Firefly* luciferase mRNA translation while two tested G-rich ODNs/ONs did not show translation repression activity (**Figure 1B**), clearly indicating the variability in the mechanism of action of G-rich ODNs/ONs.

**Figure 1.**
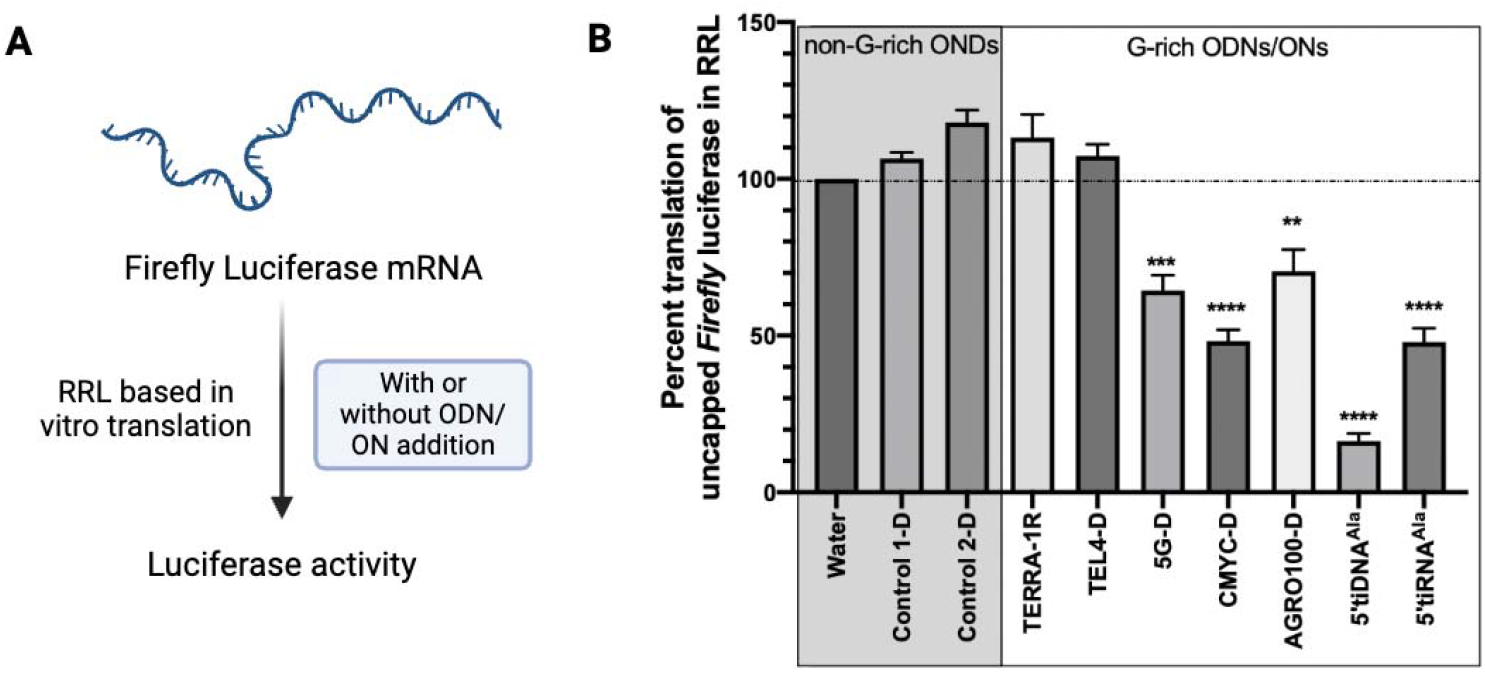
Some but not all G-rich ODNs/ONs repress reporter mRNA translation in RRL. **A**.Schematic of the experimental setup to study the impact of G-rich ODNs/ONs on translation and **B**. Several G-rich ODNs repressed the luciferase mRNA translation.

To get a clearer answer on what makes certain G-rich ODNs more potent in translation repression compared to others, we decided to expand our candidate G-rich ODN pool to include wide variety of sequence and length composition. In our expanded pool, we picked several G-rich ODN candidates which have already been studied in the past for their different structural and functional purposes **(Figure 2A and Supplementary Table 1)**. Our expanded pool includes G-rich ODNs with wide variations, for example: G-rich ODNs those start with the G-stretches in the 5’ end and those that do not; ODNs that have several two G stretches separated by various non-G nucleotides; ODNs that have several three and four G stretches. Because mRNA translation inhibition is one of the key established molecular events of several G-rich motif containing oligos, we performed several IVT experiments in the presence of extended pool of G-rich ODNs (**Figure 2**).

**Figure 2.**
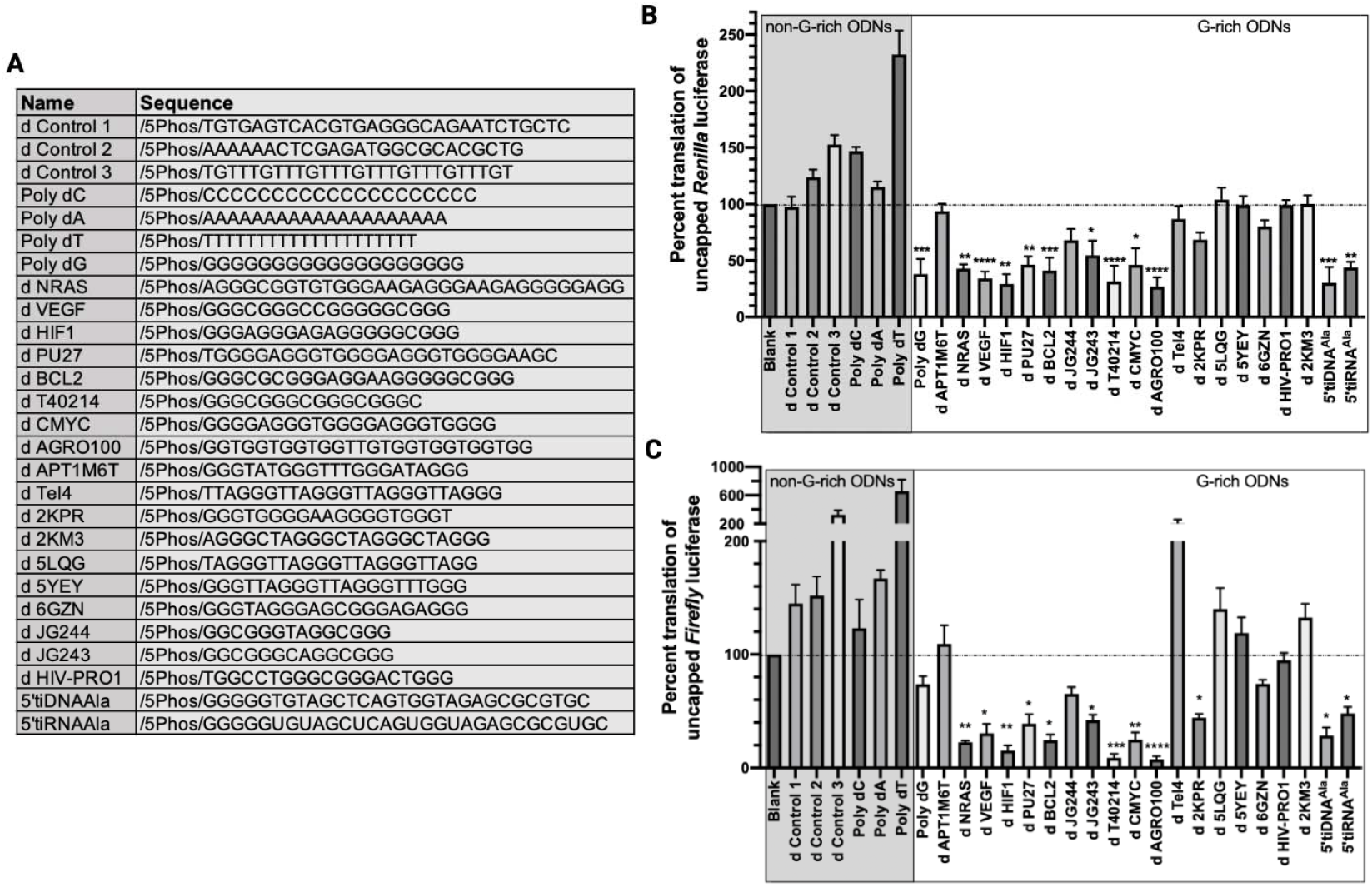
Different G-rich ODNs have different impact on human embryonic kidney derived (HEK293) cell lysate-based translation of reporter mRNAs. **A**.G-rich and non-G-rich control Oligonucleotides used in the study (further details on these oligos is available in Supplementary Materials Table 1). **B**. Relative translation efficiency of Renilla luciferase mRNA in HEK293 cells derived lysate in presence of different ODNs. **C**. Relative translation efficiency of Firefly luciferase mRNA in HEK293 cells derived lysate in presence of different ODNs. The bars represent the mean □ + □/-SEM of three independent experiments, statistical significance was tested by One-way ANOVA using water treatment as a blank control, Multiple comparisons, Dunnet correction, and p-values indicated only where there is a significant repression. * p<0.5, ** p<0.05, *** p<0.0005, and **** p<0.00005.

Despite having some advantages such as high mRNA translation efficiency and reduced preparation time, RRL-based *in vitro* translation systems have several limitations that could result in bias of the resultant outcomes^28^. For this reason and in addition, we run our IVT experiments in custom-made HEK293 cells derived cell-lysate to study the impact of different ODNs in eukaryotic translation. We prepared *in vitro* translation lysates from HEK293 cells and analyzed the ability of G-rich ODNs to inhibit the translation of different mRNA luciferase reporters in these systems. We extended our translation repression experiments for several mRNA types: uncapped *Firefly* luciferase mRNA, capped *Firefly* luciferase mRNA, uncapped *Renilla* luciferase mRNA, and capped *Renilla* luciferase mRNA. As presented in **Figure 2B-C and Supplementary Figure 1-2**, our data demonstrated that as in the case of RRL based translation, several G-rich ODNs repressed uncapped *Firefly* luciferase mRNA translation in HEK293 cell lysates. Noticeably, from our expanded pool of G-rich ODNs, several more G-rich ODN candidates seemed to be unable to repress reporter mRNA translation in any of the systems we tested.

Previously, engineered G-rich ODNs such as JG243 and JG244 were shown to inhibit HIF-1α in human cancer cells which selectively inhibited the expression of HIF-1-regulated proteins but did not impact the expression of HIF-1 non-target genes^3^. In our assays, these oligos repressed reporter mRNA translation (**Figure 2A-B**).

Importantly, the behavior of G-rich ODNs remain mostly similar on all the translation systems we tested. Specifically, G-rich ODNs, such as VEGF, AGRO100, CMYC, T40214, etc seemed to repress translation as we previously reported for 5’TOG motif containing 5’tiRNA^Ala^ and its DNA analog, 5’tiDNA^Ala 7^. However, the other class of G-rich ODNs came to our surprise as a non-translation repressing G-rich ODNs, such as TEL4, 2KM3, 5LQG, 5YEY, etc. While analyzing the sequence composition, it was obvious to us that it was not merely a G-richness of the sequences or the presence of poly G-track on the 5’end that imparts their translation repression activity.

### Bioactive ODNs form topologically different G4s than the inactive ODNs

G-rich nucleic acid sequences can fold into four stranded secondary structures called G-quadruplexes (G4s) via the stacking of square planar guanine assemblies in monovalent cation-assisted manner (**Figure 3A**)^29^. Depending upon the nature of the sequence and their surroundings, G-rich nucleic acid sequences can also fold into different G4 topologies^30^. Both Watson-Crick and Hoogsteen sides of G base are involved in H-bonding in the formation of G4s^29^. While almost all reported RNA G4s and many DNA G4s have parallel topologies, a conformational switch around glycosidic bond resulting in *syn* conformation (comparison to sterically favorable *anti* conformation) allows the formation of antiparallel loops and hence anti-parallel G4s in DNA^31^. Since most of the G-rich ODNs had potential to form G4s but they possessed different bioactivity, we reasoned that their difference in G4 topology could be contributing to their function. To test this hypothesis, we monitored the G4 folding behavior of G-rich ODNs using circular dichroism (CD) spectroscopy. In CD, a cation responsive (higher intensity at G4 permissive K^+^ or Na^+^ environment and lower at G4 non-permissive Li^+^ environment) positive peak at ∼263 nm and a trough at ∼240 nm is an indicative of a parallel G4 (p-G4) topology while a peak at ∼295 nm indicates an anti-parallel G4 (ap-G4) topology. To test the G4 topologies shown by G-rich ODNs, they were folded in different buffers at 10 μM oligonucleotide concentration and analyzed using CD spectroscopy. Interestingly, all the bioactive (translation repressing) G4 oligos show parallel topology (**Figure 3B** top panel, three representative examples are shown, AGRO100, VEGF, and CMYC; and **supplementary Figure 3**) while those not showing bioactivity showed antiparallel topologies (**Figure 3B** bottom panel, three representative examples are shown, 2KM3, 5LQG, and 5YEY; and **supplementary Figure 3**).

**Figure 3.**
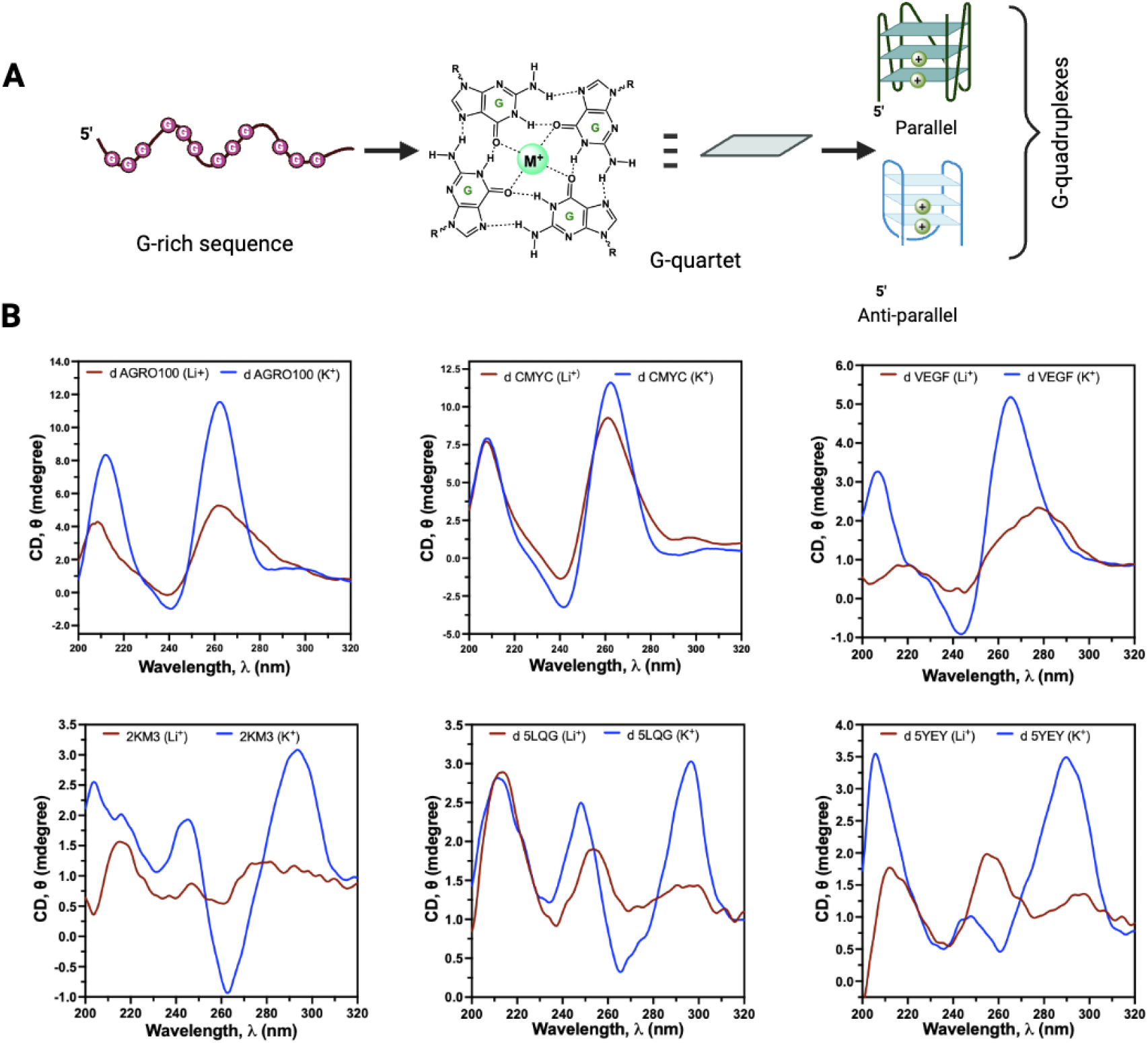
G-rich ODNs can fold into topologically different G-quadruplexes (G4s), A. A schematic demonstrating G-rich sequence can fold into parallel and anti-parallel G-quadruplex structures (among others) via the stacking of square-planar G-quartet arrangements, and B. CD spectra to demonstrate translation-repressing G-rich ODNs fold into parallel G4s (top panel) while inactive G4s fold into antiparallel G4 conformations.

### Parallel to anti-parallel switch of G4s (without changing the base composition) recues G4 ODNs bioactivity

Next, we asked whether the functional differences of p-G4 vs ap-G4 ODNs are truly due to their structural and topological variations and not because of the differences in the sequence composition. To test this, we modified translationally inactive ap-G4 ODNs to their RNA counterparts. Since the presence of 2’-OH group enforces parallel topology in RNA G4s and almost all reported RNA G4s form pG4s, we reasoned that DNA to RNA conversion of G-rich ODNs should lead to parallel topology G4s (p-rG4s). If the difference in the bioactivities was indeed a result of their topological variations, ap-G4 to p-rG4 interchange should result in the reversal of G4 function–hence transforming inactive ap-G4s to active p-rG4s.

To test this hypothesis, we synthesized RNA counterparts of three ap-G4s (namely, 2KM3, 5YEY and 2JQG). First, we confirmed their topological variations using CD spectroscopy, where we noticed expected topological switch as demonstrated in **Figure 4A**, DNA→RNA conversion of the ap-G4 ODNs resulted in p-G4 ONs as evidenced by the CD peak switch from ∼295 nm for an ap-G4 to ∼263 nm for a p-G4. Next, we asked whether the topological switch in the G4 conformation is reflected in their bioactivity. We performed IVT experiments as described before. As demonstrated in **Figure 4B-C**, ODN to ON conversion of G-rich sequences not only changes their topologies, but also inverses their biological functions-i.e. translationally inactive ap-G4s are transitioned to translation repressing pG4s. Importantly, topology dependent reversal of the action of G-rich ODNs is evident in both capped and uncapped luciferase mRNA translation.

**Figure 4.**
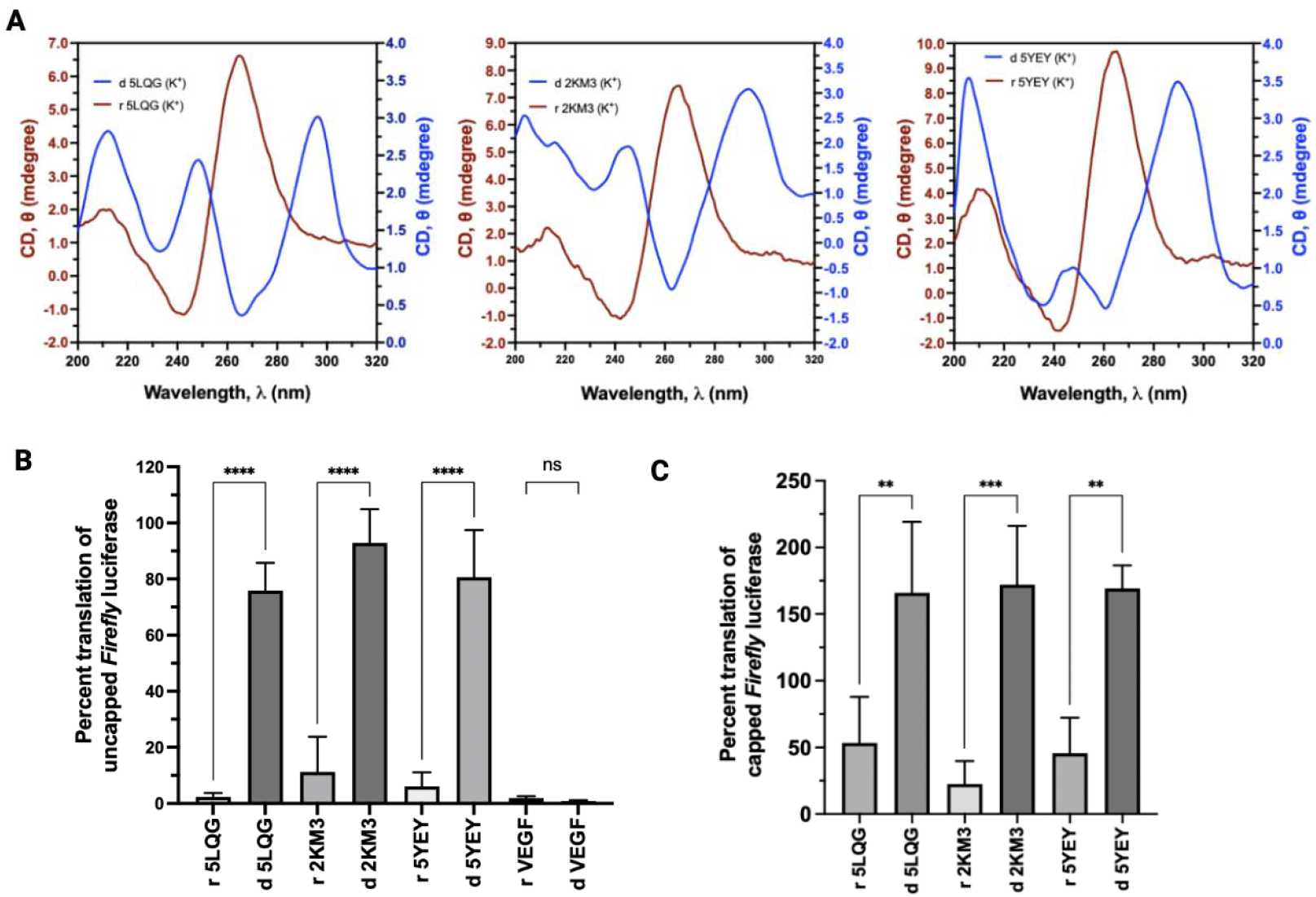
Conversion of non-bioactive G-rich ODNs to G-rich ONs not only changes their topologies but also rescued corresponding bioactivities. **A**.CD spectra to demonstrate the change in topology of anti-parallel G4 forming G-rich ODNs when they were converted into corresponding ONs (DNA→RNA). The bars represent the mean□+ □/-SEM of three independent experiments, statistical significance was tested by One-way ANOVA, Multiple comparisons, Dunnet correction, ** p<0.05, *** p<0.0005, and **** p<0.00005.

### Only parallel topology G4s directly interact with eIF4G1 via its HEAT1 domain

We previously demonstrated that G4 forming RNAs (parallel G4s) can directly interact with eukaryotic translation initiation factor 4G 1 (eIF4G1) via its first HEAT domain (HEAT1)^18^. Because of the observation that only parallel G4s can repress eukaryotic mRNA translation in different lysate systems, we reasoned that antiparallel G4s might interact with eIF4G1-HEAT1 differently. To test this hypothesis, we performed *in vitro* binding experiments of different G-rich ODNs with recombinant eIF4G1-HEAT1 and analyzed the resultant binding using a non-denaturing gel shift assay. We picked AGRO100 as a p-G4 candidate, and 5-YEY as n ap-G4 candidate for these binding experiments. Interestingly we observed that only the parallel G4 (d AGRO100) bind to eIF4G-HEAT1 with significantly low Kd (724.8 nM) (**Figure 5A** and **D**) while the antiparallel G4 (d 5YEY) did not bind to eIF4G-HEAT1 at the given concentration as evidenced by a weaker gel-shit and flatter binding curve for the later (**Figure 5B** and **D**). On the other hand, when we noticed an increase in the binding affinity of parallel G4 forming (r 5YEY) with the eIF4G1-HEAT1 as evidenced by the rescued binding in **Figure 5C-D**.

**Figure 5.**
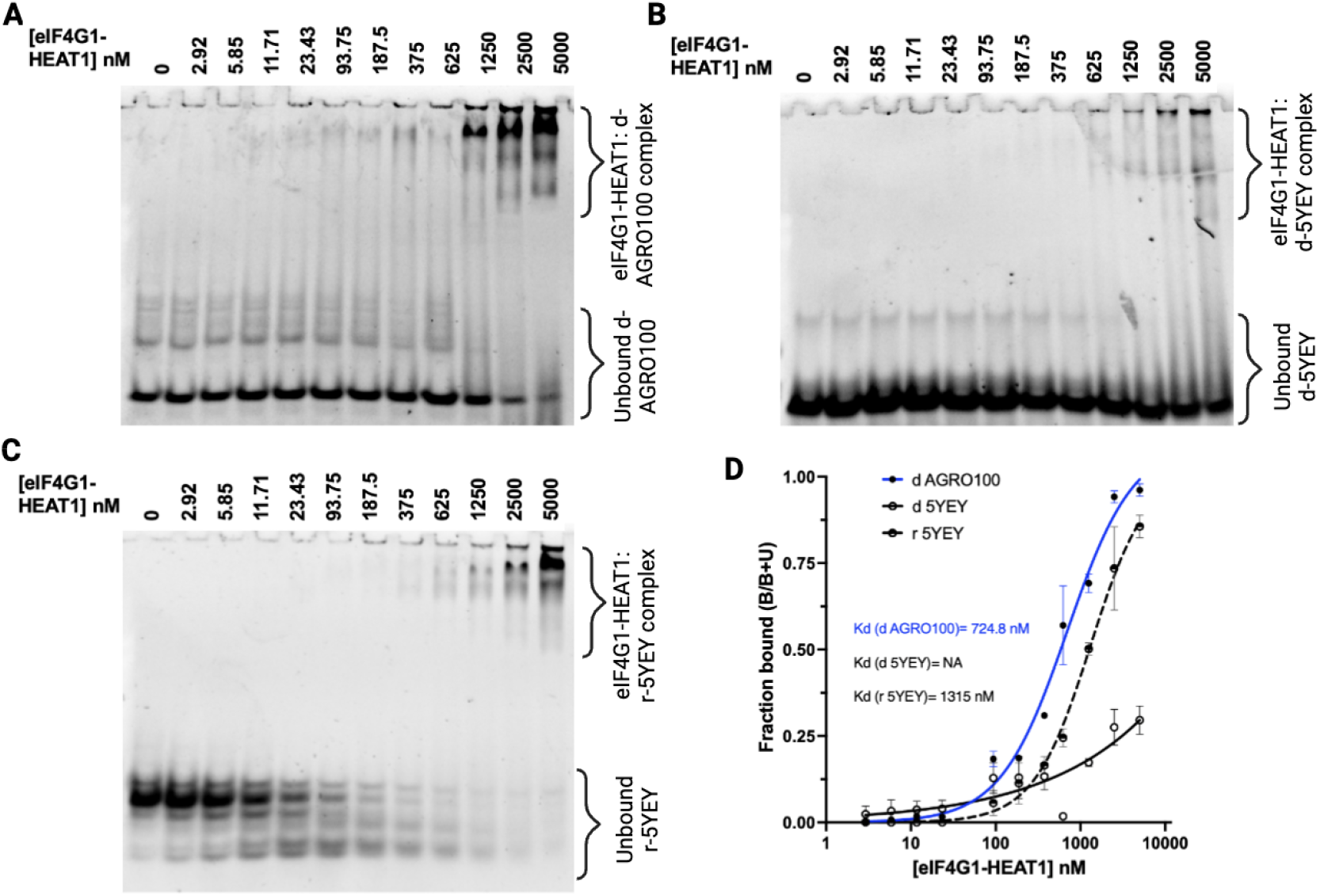
Anti-parallel G4 topology is detrimental in G4-eIF4G1 interaction. A. Parallel G4 forming AGRO100 binds with HEAT1 domain of eIF4G1 in concentration dependent manner. B. However, antiparallel G4 forming 5YEY does not bind significantly with HEAT1. C. The modification of d 5YEY to r 5YEY rescues 5YEY-HEAT1 binding suggesting the need of parallel topology for the interaction of G4s with eiF4G1. D. Nonlinear regression curve fitting for respective gel-shifts (A-C) showed a tight binding between parallel d AGRO100 and HEAT1 (blue curve) with a Kd of 724.8 nM. As evidenced from the gel-shits, there is no significant binding of anti-parallel G4 with HEAT1(solid black curve). Importantly, r 5YEY binds with HEAT1 with a Kd of 1.31 μM.

## DISCUSSION

Despite a plethora of studies investigating the potential of G-rich ODNs/ONs for their therapeutic applications^4-6,19^, we lack a clear mechanistic, biological and structural understanding of how some G-rich ODNs/ONs behave differently than the others. In this report, using a series of biochemical, biophysical, and cell biology techniques, we solved a part of the G-rich ODNs bioactivity puzzle. Our data made it clear that the translation inhibitory behavior of G-rich ODNs/ONs is mediated through their ability to fold into a particular topology G4 structure.

G-rich ODNs have been studied as molecules affecting cellular physiology^4^. They can be cytoprotective as well as cytotoxic based upon their sequence and length identities. While several of cytoeffective ODNs reported in the literature are G4 forming ODNs, how they impart the biological outcome in trans is unclear. The impact of G4s formed within both DNA and RNA molecules (e.g., telomeric sequences, 5’- and 3’-untranslated regions of mRNAs) in *cis* has been widely studied and such structures by themselves or in association with G4-binding proteins (G4BPs) were postulated to regulate almost every aspect of nucleic acids biogenesis, metabolism and gene regulation^32-34^. However, it has been overlooked in the field why some *trans*-acting G4 ODNs/ONs are cytoxic and some other are cytoprotective? By studying a series of G-rich ODNs that differ in their sequence length, composition, and topologies for their ability to impact eukaryotic translation in different systems, we discovered that only the parallel topology G4-ODNs can inhibit translation, one of biological outcomes where G4s are traditionally studied.

Because wepreviously showed that the mechanism of mRNA translation inhibition mediated by the RNA G4s involved the displacement of protein factors engaged in translation initiation^7,18,25^, we reasoned that the bioactive ODN G4s may also directly interact with the components of the mammalian translational machinery and/or translational regulators. This motivated us to ask what are the crucial sequence or structural determinants that make G-rich ODNs potent translation modulators.

G4 structures have been previously shown to interact with several G4BPs thereby influencing the downstream biology^26,34,35^. We and others previously discovered that, once transfected in the cells, G4 ODNs/ONs can interact with cellular stress response proteins such as G3BP1, YB1X, TIA1, and eIF4G1; and can induce SG assembly formation and translation repression^7,18,23,25,26,36^. However, our findings in this report revealed that translation suppression ability of G4s is not universal, rather it is topology dependent. Only parallel topology G4s can directly interact with the HEAT1 domain of eIF4G1 protein thereby dissembling the eucaryotic translation initiation complex. Contrarily, the anti-parallel topology G4s do not interact directly with eIF4G1 and hence do not show translation inhibitory effects.

## CONCLUSION

In sum, we identified the molecular mechanism by which G-rich ODNs can bestow different biological effects. We discovered that G-quadruplexes can have totally distinct biological outcomes depending upon their topological differences. Furthermore, G4 binding to HEAT1 domain of eIF4G1 can clearly distinguish between a parallel vs anti-parallel topology G4 resulting in completely different functional consequences. Our findings in this study widen our understanding of mechanistic differences created by topological variabilities within the G-quadruplex forming therapeutic oligodeoxyribo-nucleotides.

## Author Contributions

PK, SML and PI developed the idea, PK designed the experiments, performed most of the experiments, and analyzed the data, PK and PI supervised the project; NB performed IVT experiments, SZ prepared general reagents; PA provided funding support, participated in discussion; PI provided the funding support, supervised the project; SML participated in the discussion, PK and PI wrote the manuscript. All authors reviewed the manuscript and agreed on the final format.

## Notes

The authors declare no competing financial interest.

## Supporting Information

Supporting Information is available as a separate attachment.

## Acknowledgements

We are thankful to all Anderson-Ivanov lab members for their valuable input.

## Entry for the Table of Contents

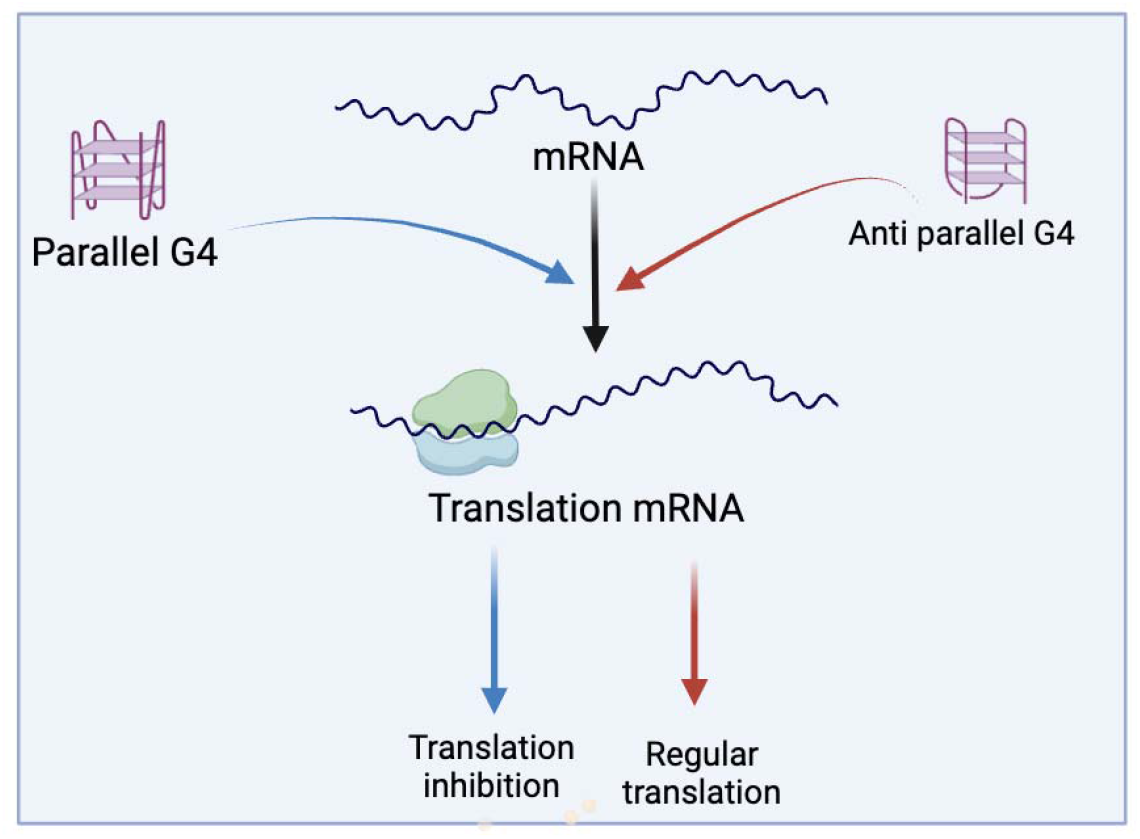

**TOC**. Therapeutic G-quadruplex harboring oligo deoxyribonucleotides can impart totally different regulatory functions depending upon their topologies. Only parallel topology G4s directly interact with translation machinery and can inhibit protein synthesis.

