## Supplementary Info for "G-quadruplex topologies determine the functional outcome of guanine-rich bioactive oligonucleotides"

**Materials and Methods**

**ODNs/ONs**

All the ODNs and ONs were purchased from IDT. Stock solutions were made up to a concentration of 100 μM in nuclease-free water and stored at −80°C. The oligonucleotides used are listed in Figure 2A and supplementary materials Table 1. 3’-end biotinylated oligonucleotides had the identical sequence in which the 3’-end is biotinylated for the pull-down applications.

**Cell culture**

Human osteosarcoma (U2OS) cells and human embryonic kidney derived (HEK293) cells were maintained in DMEM medium (Gibco) containing 10% fetal bovine serum (FBS) with 1% penicillin–streptomycin. Cells were either used for translation lysate preparation (as described below) or of the study of stress granule formation. Cells were transfected with 250 pmol of indicated oligonucleotide for 6 h using Lipofectamine 2000 (Invitrogen). Cells were manually scored for SGs using fluorescence microscopy with eIF4G1 and G3BP1 as SG markers. Only cells with granules co-staining for both markers were considered SGs, and a minimum of 3 SGs per cell were required to score positive.

**Reporter mRNA preparation**

*Firefly luciferase* mRNA was bought from Promega (Cat# L4561) and capped using cell script m7G capping kit (Cat# C-SCCS1710) as per manufacturer’s protocol. *Renilla luciferase* reporter mRNA was transcribed in vitro using Promega Ribomax T7 transcription kit (Cat# P1300) using SacII digested pHRL control plasmid.

**Preparation of translation competent mammalian extract**

After reaching 80-90% confluency, HEK293 cells were detached via trypsinization at 37°C for 2min. Equal volume of DMEM medium was added to stop the trypsinization reaction and cells were pelleted at 5000rpm for 2min. The media was removed, and cells were suspended in 1ml of hypotonic lysis buffer (10mM HEPES pH7, 10mM potassium acetate, 0.5mM magnesium acetate, 5mM DTT and one tablet of mini EDTA free protease inhibitor tab from Roche). The cells were transferred to a 1.5ml microcentrifuge tube and centrifuged at 1000 RCF at 4°C for 1min. The supernatant was removed, and the cells were resuspended in chilled lysis buffer (volume of lysis buffer is same as volume of packed cells). Cells were tumbled at 4°C for 1hr followed by homogenization by passing through a 1ml syringe 27G needle for 10-12 times. The resulting lysate was centrifuged at 14000 RCF for 1 min and 40ul of supernatant was aliquoted in 1.5ml microcentrifuge tubes. The tubes were snap frozen in dry ice followed by immediate storage at -80°C.

***In vitro* translation**

*In vitro* translation reaction in RRL was performed as per manufacturer’s protocol with some changes. Briefly, a 10µl reaction was set up with 10pm of respective oligos incubated with 7µL of RRL, 0.1µL of any two amino acid mix, 50-100ng of reporter mRNA and 10 U of RNase inhibitor at 30°C for 30 min. 2 µL of the reaction was mixed with 48 µL of luciferase assay substrate in 96-well corning flat white plate and luminescence were measured in Promega Glomax Explorer Machine (Cat ID: GM3500). *In vitro* translation reaction in mammalian extracts were performed as described in previous report24. Briefly, a 10 µL reaction was set up with 10 pM of respective ODNs with 100 ng of reporter mRNA, 1.25 mM of magnesium acetate, 150 mM potassium acetate, 1X translation buffer (1.6mM HEPES pH7, 2mM creatine phosphate, 0.01 µg/mL creatine kinase, 10 µM spermidine and 10 µM L-amino acid mix, cat ID # L4461), 1 unit of RNase inhibitor (Promega, cat ID# N251B) and micrococcal nuclease (Thermo Fisher, cat ID# 88216) treated mammalian extract. The tubes were then incubated at 37°C for 1 h followed by taking luminescence reading. Depending upon the reporter mRNA used, 20-50% of the reaction was mixed with either Firefly or Renilla and luminescence was measured using Promega Glomax Explorer Machine.

**ODNs/ONs folding**

ODNs/ONs were folded in 150 mM KCl or NaCl or LiCl in T10E0.1 buffer (10 mM Tris-HCl pH 7.4 and 0.1 mM EDTA) by heating at 95 °C for 5 min and then slowly cooling to room temperature.

**CD spectroscopy**

JASCO J815 spectropolarimeter was used to collect CD spectra. The oligonucleotides were dissolved in 150 mM K+ in T10E0.1 buffer. Quartz cuvettes (1 mm path length) were used with sample volumes of 200 μL to achieve a sample concentration of 5 μM. Spectra were collected in the range between 220 and 320 nm at 20 °C from three scans, and a buffer baseline was subtracted from each spectrum. CD was expressed as the difference in the molar absorption of the right-handed and left-handed circularly polarized light. An increased peak intensity of the oligo under K^+^ environment at 260-265 nm and a trough at 240 nm, which shows a reduced peak intensity under Li^+^ environment suggests the formation of a parallel topology G4, whereas a cation responsive peak at around 295 nm indicates an anti-parallel topology.

**Electrophoretic mobility shift assays**

Synthetic ODNs/ ONs and proteins were mixed in 10 mM sodium phosphate (pH 7.5), 300 mM NaCl, 2 mM β-mercaptoethanol, 5 mM MgCl_2_, 5% glycerol, 1 mM ATP and incubated at room temperature for 15 min before loading onto 20% acrylamide gel (29:1 Acrylamide:Bis-acrylamide, 0.5X TBE). The gels were stained with SYBR Gold and scanned.

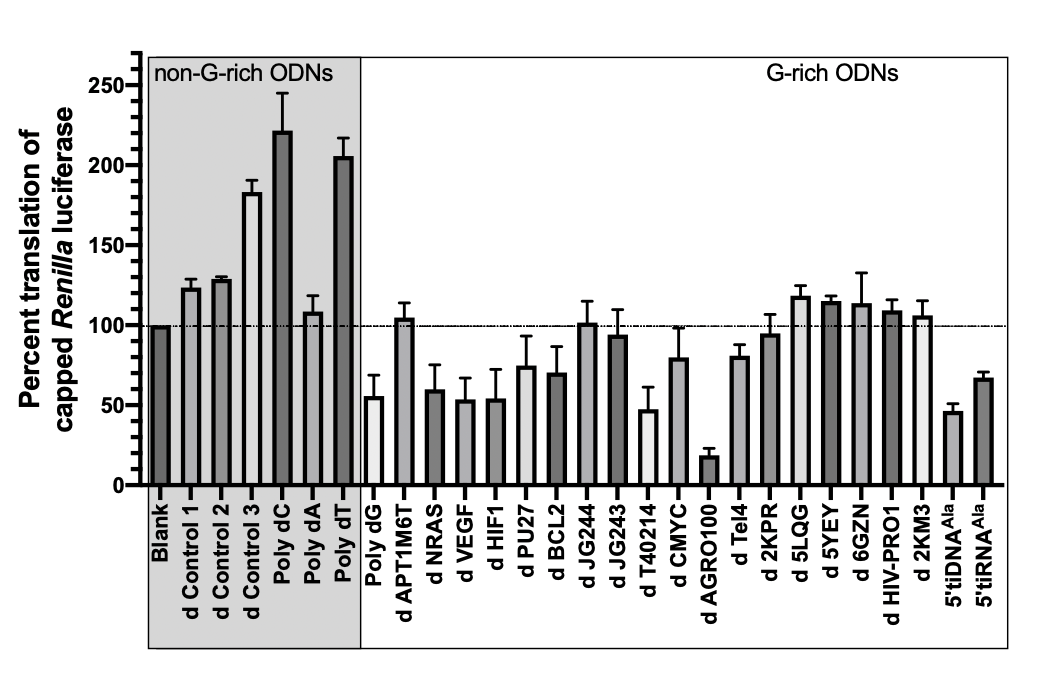

**Supplementary Figure 1.** Effect of G rich ODNs on capped *Renilla luciferase* translation.

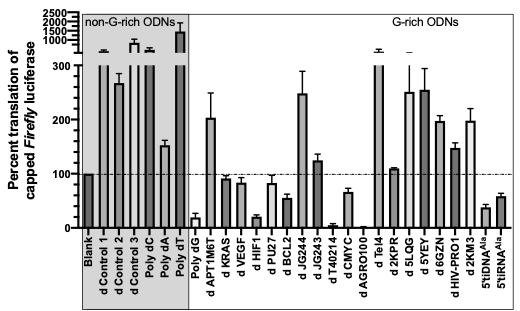

**Supplementary Figure 2.** Effect of G rich ODNs on capped *Firefly luciferase* translation.

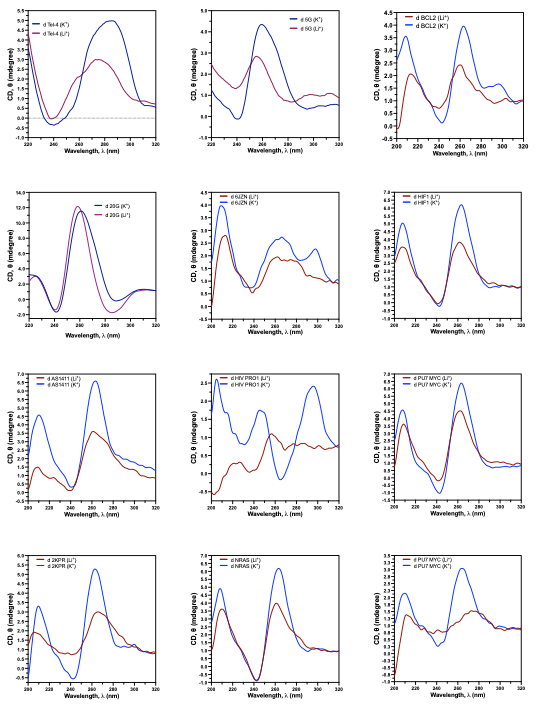
**Supplementary Figure 3.** CD spectra of additional oligos used in this study.
